# APOL1 is not expressed in proximal tubules and is not filtered

**DOI:** 10.1101/623918

**Authors:** Natalya A. Blessing, Zhenzhen Wu, Sethu Madhavan, Myung K. Shin, Maarten Hoek, John R. Sedor, John F. O’Toole, Leslie A. Bruggeman

**Affiliations:** Rammelkamp Center for Education and Research, MetroHealth Medical Center, Case Western Reserve University School of Medicine, Cleveland, OH 44195; Departments of Inflammation & Immunity and Nephrology, Cleveland Clinic, Case Western Reserve University School of Medicine, Cleveland, OH 44195; Department of Physiology & Biophysics, Case Western Reserve University School of Medicine, Cleveland, OH 44195; MRL, Merck & Company, Inc., Kenilworth, NJ 07033.; Maze Therapeutics, South San Francisco, CA 94080

## Abstract

The kidney expression pattern of APOL1 was examined using both protein and mRNA *in situ* methods on *APOL1* bacterial artificial chromosome transgenic mice, with and without proteinuria. APOL1 was detected in podocytes and endothelial cells of the kidney, but was not expressed in tubular epithelia, nor was plasma APOL1 protein filtered and reabsorbed by the proximal tubule. APOL1 expression in podocytes and endothelia should remain the focus for mechanistic studies of APOL1-mediated pathogenesis.

## Introduction

The mechanism of pathogenesis associated with African *APOL1* polymorphisms and risk for non-diabetic chronic kidney disease (CKD) is not fully understood. Prior studies have minimized a causal role for the constitutively circulating APOL1 protein (1–3), thus efforts to understand kidney pathogenesis have focused on APOL1 expressed in renal cells. The *APOL1* kidney expression pattern remains unclear with published discrepancies between immunohistochemistry and mRNA *in situ* hybridization results, most notably the abundant APOL1 protein observed in proximal tubule cells (4–6). In addition, since APOL1 is constitutively present in blood, it is unclear if APOL1 is filtered, especially in the setting of proteinuria, which could result in APOL1 protein reabsorption by the proximal tubule.

APOL1 in circulation is bound to either a 500kDa HDL_3_ particle, known as trypanolytic factor 1, or to a 1000kDa lipid-poor IgM complex, known as trypanolytic factor 2 (7–9). The proteins produced by the two CKD-associated *APOL1* variant alleles, G1 and G2, bind these high molecular weight complexes similar to the common allele G0 (10). Although the APOL1 protein (42.5kDa) is small enough to pass the glomerular filtration barrier size restriction limit, it is not known to circulate independent of these high molecular weight complexes (11). However, lipoproteins and other components of HDLs can be filtered (12), and in the setting of proteinuria, larger molecular weight proteins may appear in filtrate, exposing tubular epithelia to serum proteins not normally filtered. It is unclear whether APOL1 or APOL1-containing complexes may be filtered in the setting of proteinuria.

## Results

Three transgenic mouse lines expressing a human bacterial artificial chromosome (BAC) encompassing the entire *APOL1* genomic region for either the G0, G1, or G2 alleles have been described (13, 14). These mice express *APOL1* under native gene regulatory regions and appear to replicate the gene expression pattern observed in humans. APOL1 protein was abundant in podocytes and also was present in endothelial cells of peritubular capillaries, glomerular capillaries, and arteries (Fig. 1A,B). This pattern in the BAC-APOL1 mice was consistent with our prior reports examining human kidney tissue (6). Expression in podocytes and endothelia was confirmed with mRNA *in situ* hybridizations, however, no APOL1 protein or mRNA was detected in the proximal tubule or any other nephron segment (Fig. 1C). Lack of tubule expression is consistent with our prior mRNA *in situ* hybridization results from human kidney which also did not find evidence of *APOL1* expression in tubules (5). Also similar to humans, the BAC-APOL1 transgenic mice had abundant APOL1 in serum (Fig. 1E). A prior study of human liver transplant recipients, whose *APOL1* genotypes did not match the *APOL1* genotype of their allografts, established that circulating APOL1 in humans is largely produced by the liver (15). The BAC-APOL1 transgenic mice had abundant APOL1 protein and mRNA expression in the liver, predominately in zone 3 hepatocytes immediately adjacent to central veins (**Supplemental Figure 1**). In the kidney, this circulating APOL1 protein was readily detected in blood trapped in vascular lumens (**Supplemental Figure 2**). Overall, there was no difference in the *APOL1* expression patterns between the G0, G1, or G2 expressing mice (**Supplemental Figure 2**), similar to our previous studies in human biopsies from patients with different *APOL1* genotypes (5, 6, 16).

**Figure 1.**
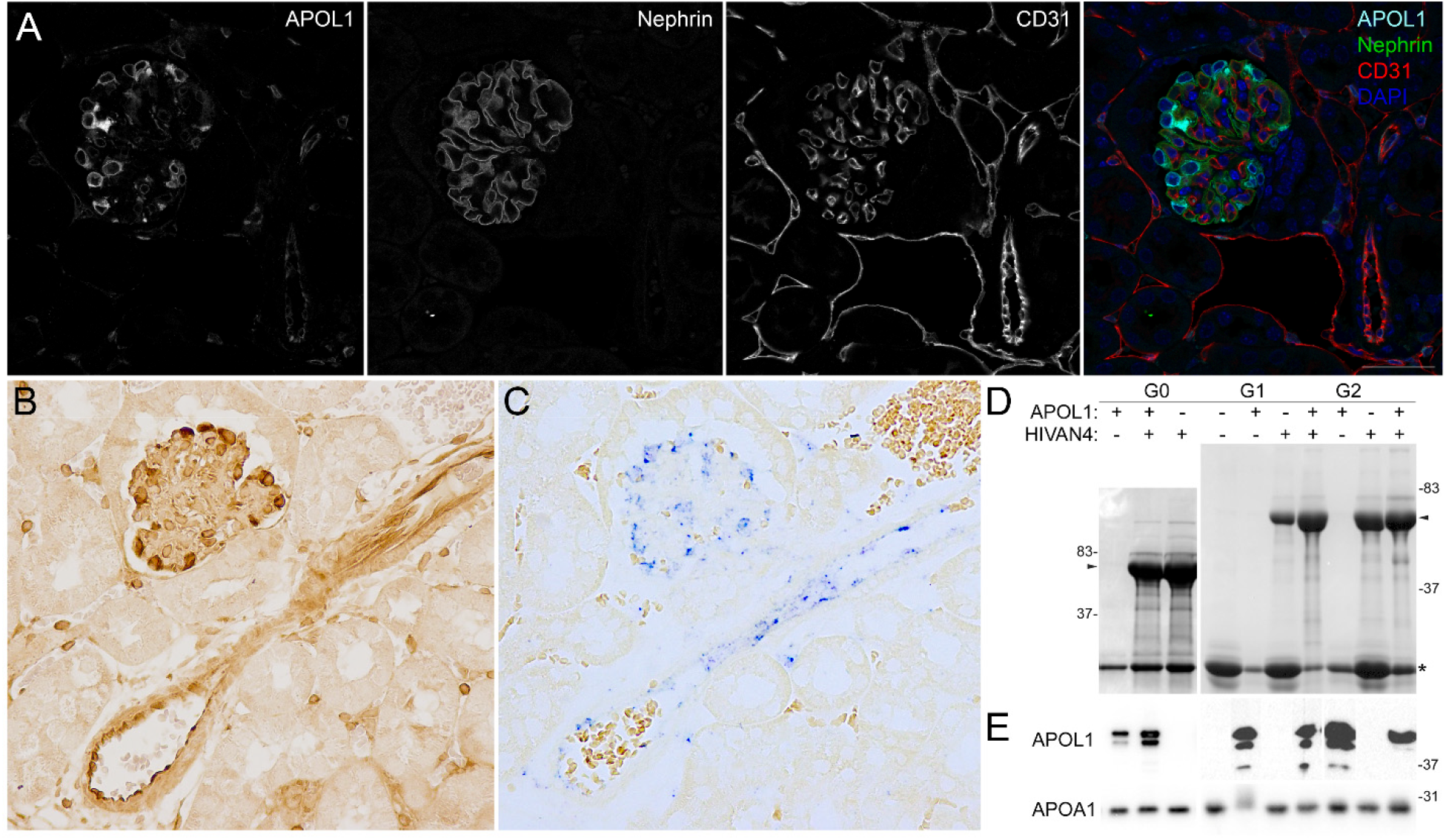
*APOL1* was expressed in BAC-APOL1 transgenic mice sera and in podocytes and endothelia. **A**. Immunofluorescent staining for APOL1, Nephrin, and CD31 in the BAC-APOL1 transgenic mouse kidney (G1 mouse is shown). Scale bar=40µm. **B-C**. Serial sections of BAC-APOL1 mouse kidney for (**B**) APOL1 protein using an immunohistochemical stain (brown, no counterstain) and for (**C**) *APOL1* mRNA expression using *in situ* hybridization (blue, no counterstain) confirming expression in kidney was restricted to podocyte and endothelium. **D**. Proteinuria in single and dual transgenic mice (coomassie stain, arrowhead marks albumin). Additional low molecular weight urinary proteins (asterisk) are a normal finding in mice. **E**. Western blot of mouse serum in the same single and dual transgenic mice showing maintenance of high levels of serum APOL1 protein in the setting of proteinuria. G0 sera were diluted 1:10, G1 and G2 sera were undiluted. APOA1 Western blot as a positive control for serum loading, all were diluted 1:10. Normal concentrations of APOL1 (42.5kDa) in human sera range from 3-30µg/ml. Normal APOA1 (28kDa) concentrations in human and mouse sera are similar and range from 0.9-2.0µg/ml.

None of the BAC-APOL1 mice spontaneously developed proteinuria (Fig. 1D), which is consistent with the original description of these mouse models (13, 14). When proteinuria was induced by interbreeding with a model of HIV-associated nephropathy (Fig. 1D), APOL1 was not evident in filtrate or within proximal tubules (Fig. 2). These mouse kidneys were also immunostained for APOA1, an apolipoprotein that is filtered, as a positive control for the vascular distribution and proximal tubule reabsorption patterns. APOA1 was readily detected in glomerular capillary lumens, and also concentrated in protein reabsorption droplets at the brush border (**Supplemental Figure 3A**). In the setting of proteinuria, APOA1-containing reabsorption droplets increased in both number and staining intensity at the proximal tubule brush border (**Supplemental Figure 3B**). A similar pattern was not observed with APOL1 (Fig. 2).

**Figure 2.**
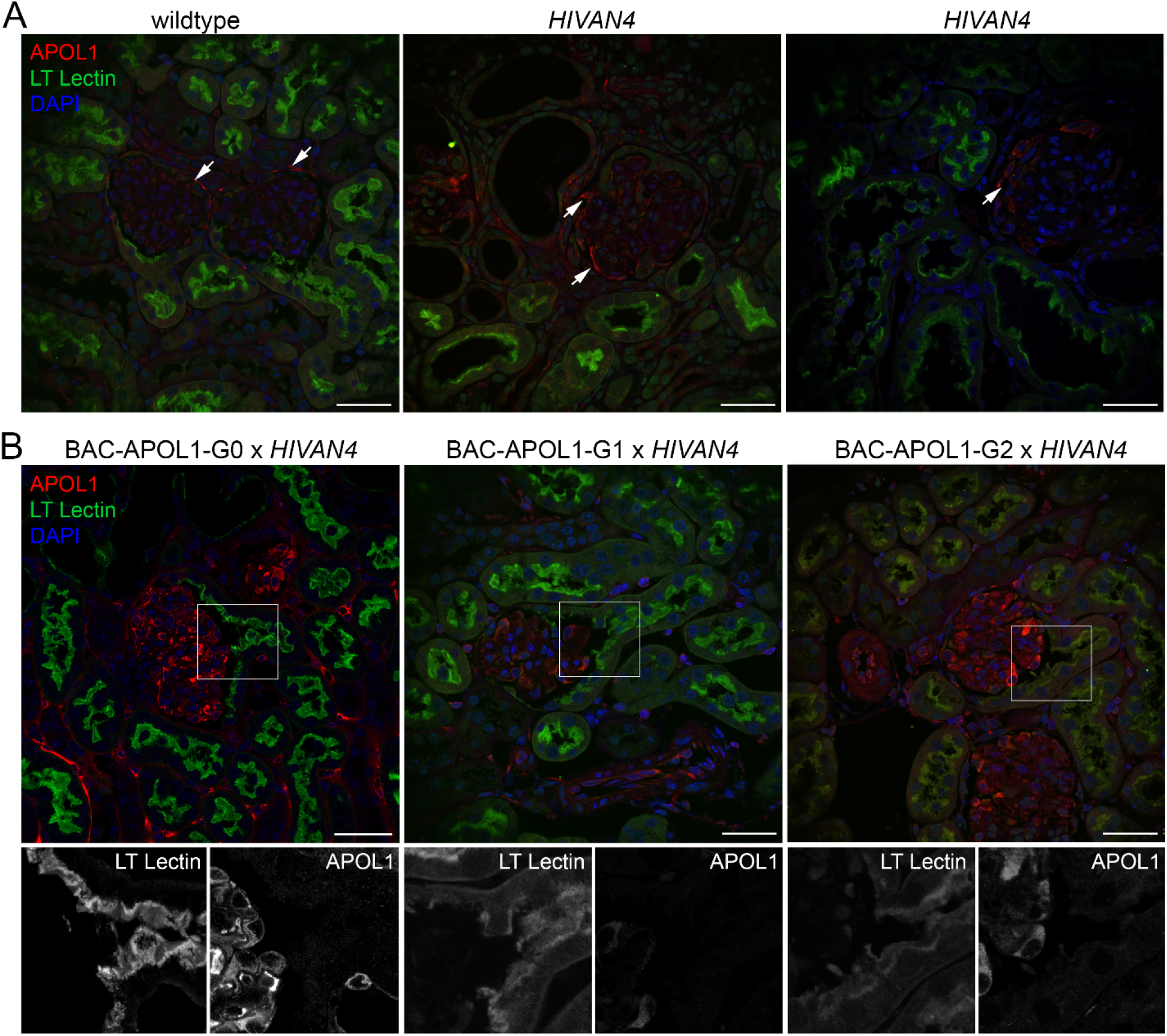
APOL1 does not appear in proximal tubules in proteinuric BAC-APOL1 transgenic mice. **A**. Control immunostaining in non-transgenic (wildtype) and proteinuric *HIVAN4* mice for APOL1 along with fluorescent lotus tetragonolobus (“LT”) lectin binding as a marker for proximal tubule cells. Since wildtype and *HIVAN4* mice do not have APOL1, the staining observed in parietal cells (arrows) is artifact. **B**. In proteinuric mice, immunofluorescence for APOL1 was not evident in proximal tubules. The boxed region is magnified below each panel along with the isolated fluorescent channels shown in black and white. Scale bar=40µm.

## Discussion

Antibody specificity has been recognized as one of the most significant challenges impacting reproducible research (17). We and others had originally reported APOL1 protein is present in proximal tubules of normal and diseased subjects (4, 6). However, our continuing work with other anti-APOL1 antibodies and subsequent studies using mRNA *in situ* hybridization indicated APOL1 was not expressed by proximal tubules (5). A remaining possibility is that APOL1 protein is filtered and reabsorbed, resulting in the appearance of APOL1 protein in proximal tubules. Using *in situ* methods that do not rely on antibodies, along with the newly available human BAC-APOL1 transgenic mice, neither APOL1 protein nor *APOL1* mRNA could be detected in any tubular epithelia. In the setting of proteinuria, APOL1 also was not filtered. In our hands, the Sigma rabbit polyclonal antibody consistently recognizes human APOL1 but has lot-to-lot differences with recognition of other proteins. The Sigma rabbit polyclonal lot used in studies here, does not replicate the strong proximal tubule staining observed in prior lots. Although monoclonal antibodies would eliminate the variation inherent in polyclonal antibody production lots, the commercial monoclonal antibodies are non-specific or have poor sensitivity for APOL1 (our experience with commercial anti-human APOL1 antibodies are summarized in **Supplemental Tables 1A-C, and Supplemental Figure 4**). The use of normal mouse kidney tissue as an irrefutable negative control is a strong quality control step in validating APOL1 antibodies and is routinely used in our laboratory.

A reproducible and consistent observation is the expression of *APOL1* in podocytes and endothelium in both the BAC-APOL1 transgenic mice in this study and in prior work with human kidney tissue. In addition, observations here indirectly support conclusions from prior studies (1, 2) that circulating APOL1 protein is unlikely to contribute to kidney disease pathogenesis as it is not the source of kidney-localized APOL1. Evaluating the contribution of podocyte- and endothelia-expressed APOL1 is a logical focus for future studies examining the mechanism for *APOL1* risk variant contributions to CKD pathogenesis.

## Short Methods

### Mouse Models

Three transgenic mouse lines expressing a 47kb human BAC encompassing the promoter and coding regions of the human *APOL1* gene for each G0, G1, or G2 alleles (BAC-APOL1-G0, BAC-APOL1-G1, or BAC-APOL1-G2) were developed at MSD Research Laboratories and have been previously described (13, 14). BAC-APOL1 transgenic lines were backcrossed to FVB/Nj for this study. The mouse model of HIVAN used to induce proteinuria was the Tg26 *HIVAN4* congenic that develops a less aggressive disease than the parental Tg26 model and has been previously described (18). Intercrossed BAC-APOL1 transgenic mice with *HIVAN4* mice were sacrificed at ~45 days of age when proteinuria was >2+ by urine dipstick (G0 n=15, G1 n=11, G2 n=9). Albuminuria was assessed by polyacrylamide gel electrophoresis of 1µl urine, followed by coomassie staining, and serum APOL1 was compared using Western blotting as previously described (19). All animal studies were with oversight and approval from the Institutional Animal Care and Use Committees of Case Western Reserve University and the Cleveland Clinic.

### Immunohistochemistry and Immunofluorescence

Formalin-fixed (overnight, 4°C), paraffin-embedded kidney tissue sections were subjected to antigen retrieval in boiling citrate buffer as previously described (6) using detection with either immunofluorescence using FITC, Cy3, or Cy5-labeled species-specific secondary antibodies or for immunohistochemistry using an avidin-biotin system (ABC kit and DAB color development, Vector Labs). Primary antibodies used include rabbit anti-human APOL1 (Sigma, HPA018885, lot E105260, 1:400 dilution), rabbit anti-mouse APOA1 (ThermoFisher, 1:500 dilution), goat anti-mouse CD31 (R&D Systems, 1:200 dilution), guinea pig anti-Nephrin (USB, 1:200). FITC-labeled lotus tetragonolobus lectin (Vector labs) was used to label proximal tubule cells as previously described (6). For commercial antibody testing, mouse kidney tissue was fixed using a variety of methods and paraffin embedded. Immunohistochemistry was performed using a variety of antigen retrieval methods and various commercial antibodies to human APOL1 (details in **Supplemental Figure 4 and Supplemental Tables 1A-C**).

### mRNA in situ hybridization

The ViewRNA ISH Tissue Assay Kit and a type 6 probe for human *APOL1* (Invitrogen) were used to assess *in situ* mRNA expression patterns on 4µm sections formalin-fixed (overnight, 4°C), paraffin-embedded mouse kidney tissue following kit instructions. Pretreatments used were 20 minute boiling time and 20 minute protease digestion time, and color development was with fast blue reagents. The fast blue reagent can be visualized with either white light (Figure 1) or by fluorescence (**Supplemental Figure 1**)

## Acknowledgements

This work was supported by the National Institutes of Health grant numbers DK108329, DK095832, and DK007470. We thank Dr. William Baldwin for assistance with antibody specificity testing.

