## Supplemental data for "APOL1 is not expressed in proximal tubules and is not filtered"

Supplemental Figure 1. APOL1 expression in liver in the BAC-APOL1 transgenic mouse models.

Supplemental Figure 2. APOL1 serum and kidney expression levels in the BAC-APOL1 transgenic mouse models.

Supplemental Figure 3. Positive control immunostaining for filtered lipoproteins.

Supplemental Figure 4. Commercially-available monoclonal anti-APOL1 antibodies non-specifically react with epitopes in both normal (non-transgenic wild-type) and APOL1 transgenic mouse kidney sections.

Supplemental Table 1A. List of commercial antibodies examined.

Supplemental Table 1B. Summary of commercial anti-human APOL1 antibody testing in Western blotting.

Supplemental Table 1C. Summary of commercial anti-human APOL1 antibody testing in human kidney tissue, and in mouse kidney tissue as a negative control.

Supplemental Detailed Methods and Summary.

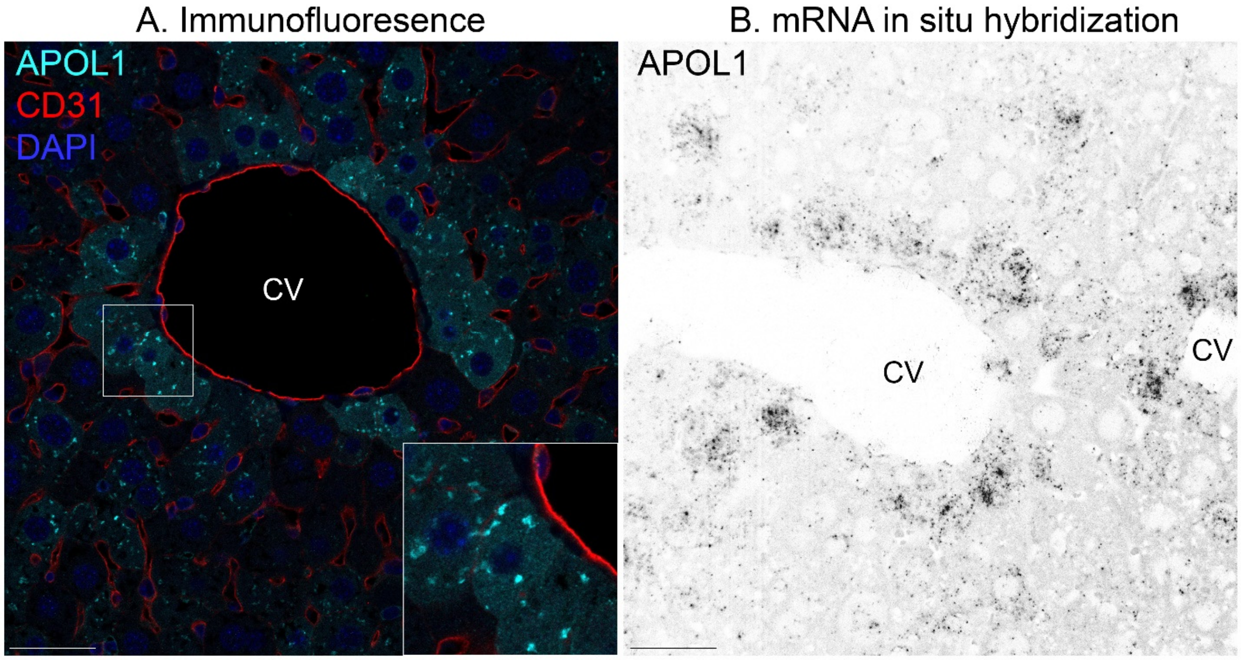


**Supplemental Figure 1. APOL1 expression in liver in the BAC-APOL1 transgenic mouse models. A**. Immunostaining for APOL1 protein with co-immunostaining for CD31 to indicate endothelial cells, and DAPI as a nuclear counterstain in a BAC-APOL1-G0 mouse. Expression was evident in hepatocytes, most strongly in zone 3 immediately adjacent to central veins ("CV"). **Inset**: APOL1 was present diffusely in the cytoplasm with more intense staining in a punctate pattern suggesting accumulation in intracellular vesicles. **B**. *APOL1* mRNA expression using *in situ* hybridization (inverted fluorescence image, no counterstain) confirming expression in liver was predominant in zone 3 hepatocytes. Scale bar=40μm.

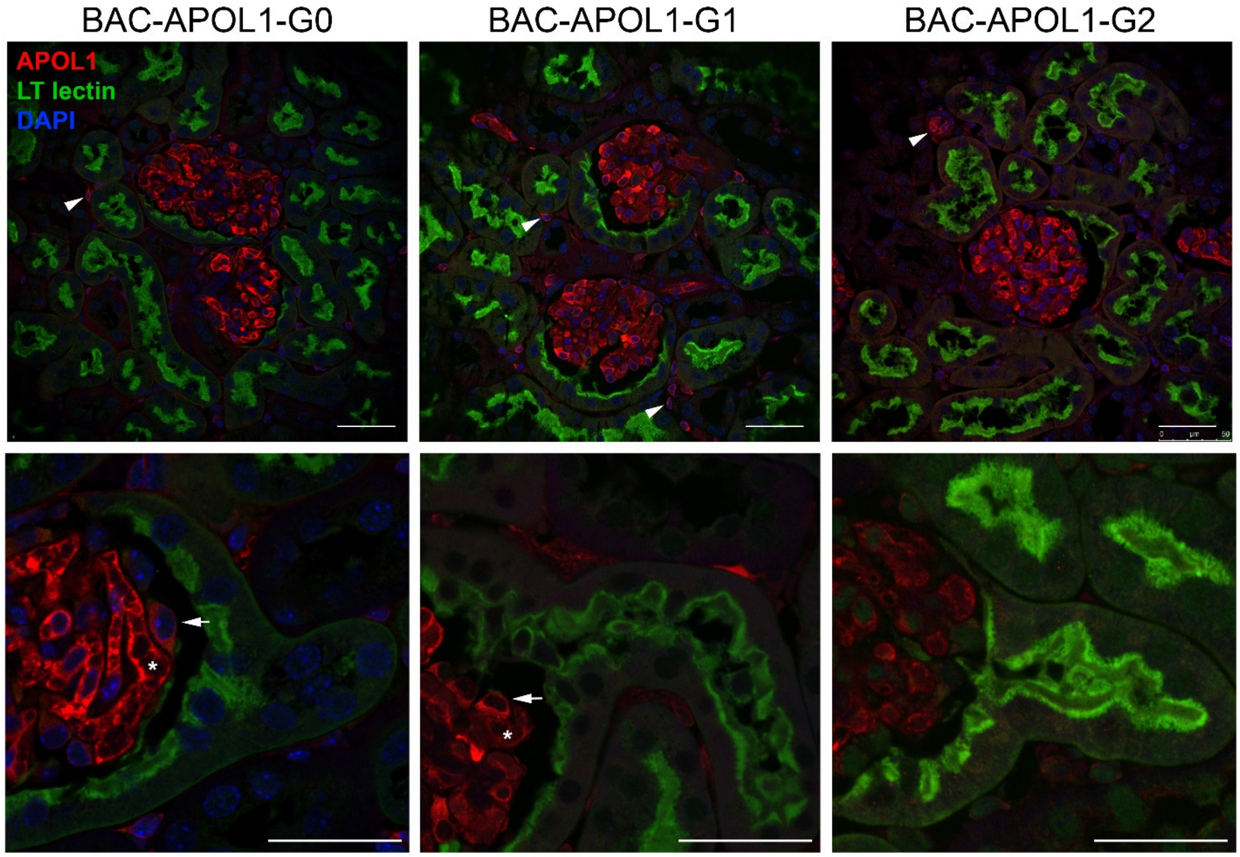


**Supplemental Figure 2. APOL1 serum and kidney expression in the BAC-APOL1 transgenic mouse models.** Immunofluorescence for APOL1, with lotus tetragonolobus ("LT") lectin staining as a marker for proximal tubule cells (arrows are podocytes, asterisks denote vascular lumen containing serum APOL1 protein, arrow heads are endothelial cells). The *APOL1* expression levels and patterns in BAC-APOL1-G0, G1, G2 mice are consistent with the original publication. There is no evidence of APOL1 protein in proximal tubules. Scale bar=40μm.

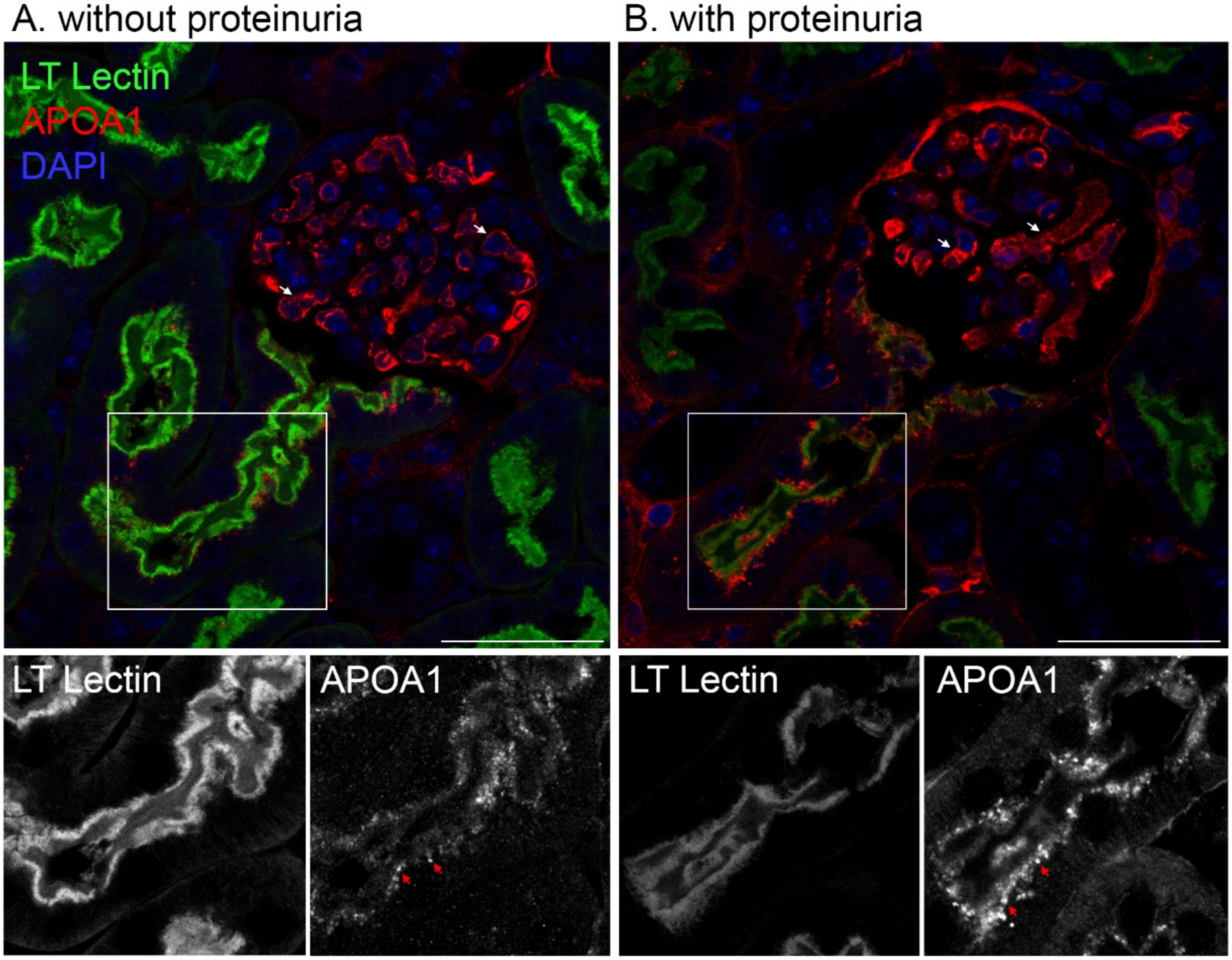


**Supplemental Figure 3. Positive control immunostaining for a filtered lipoprotein.** Immunofluorescence staining for APOA1, an apolipoprotein that is normally filtered. Sections were also stained using fluorescently-labelled lotus tetragonolobus ("LT") lectin to demarcate the proximal tubule brush border. **A**. BAC-APOL1-G0 mouse. **B**. BAC-APOL1-G0 x *HIVAN4* dual transgenic mouse with proteinuria. Below each respective color panel is the individual fluorescent channels (in black and white) of the boxed region for either LT lectin or APOA1. White arrows mark glomerular capillaries containing circulating APOA1 protein within capillary lumens, red arrows denote APOA1 in protein reabsorption droplets at the brush border of proximal tubules. With proteinuria, an enhancement in the number of APOA1-containing protein reabsorption droplets was observed. Scale bar=40μm.

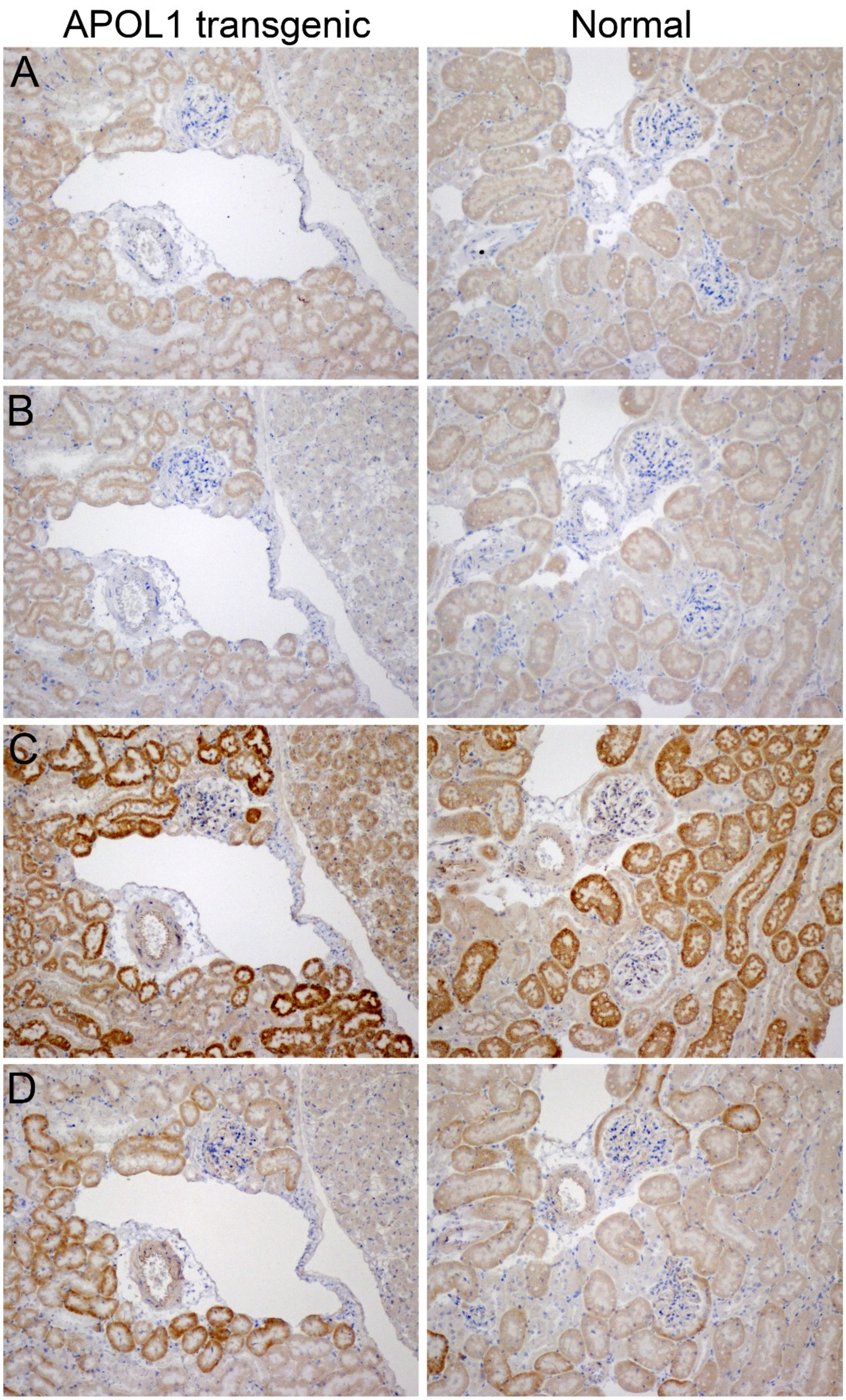


**Supplemental Figure 4.**

**Supplemental Figure 4 Legend. Commercially-available monoclonal anti-APOL1 antibodies non-specifically react with epitopes in both normal (non-transgenic wild-type) and APOL1 transgenic mouse kidney sections.** Normal mouse kidney does not express APOL1, thus any positive signal seen in normal mouse kidney is artifact. Serial sections for normal mouse and APOL1 transgenic mouse kidneys are shown with hematoxylin counterstain. Conditions for the above images were overnight formalin fixation with boiling citrate buffer antigen retrieval, using an alkaline phosphatase-conjugated secondary antibody and an avidin-biotin amplification step, generating a brown color reaction product (see Supplemental Detailed Methods). **A**. ProteinTech mouse monoclonal (66124-1-Ig). **B**. Sigma mouse monoclonal (AMAB90530). **C**. Epitomic rabbit monoclonal (n-terminal, 2840-1) **D**. Epitomics rabbit monoclonal (c-terminal, 3245-1). See **Supplemental Tables 1A-C** for a summary of other conditions tested and antibody validation results.

**Supplemental Table 1A.** Commercial anti-human APOL1 antibodies tested in our lab.

| Antibody # for Table 1B,C | Company (Catalog #) | Type | Immunogen |
|---|---|---|---|
| | Sigma (HPA018885) | **rabbit polyclonal** | aa261-385 (c-terminal) |
| 1A | lot A39110 | | |
| 1B | lot C57797 | | |
| 1C | lot E96151 | | |
| 1D | lot E114503 | | |
| 1E | lot E105900 | | |
| 1F | lot G115599 | | |
| 1G | lot E105260 | | |
| 2 | Sigma (AMAB90530)<br>also sold by other vendors | **mouse monoclonal**<br>clone CL0170 | aa261-385 (c-terminal)<br>(mapped to aa368-382) |
| 3 | Epitomics (3245-1)/Abcam (ab108315) | **rabbit monoclonal**<br>clone EPR2907 | aa305-330 (c-terminal) |
| 4 | Epitomics (2840-1)/Millipore (MABS387)<br>also sold by other vendors | **rabbit monoclonal**<br>clone EPR2906 or ID4 | aa65-90 (n-terminal) |
| 5 | ProteinTech (66124-1-lg) | **mouse monoclonal**<br>clone 1G12D11 | near full length (238aa)<br>n-terminal GST fusion |

**Supplemental Table 1B.** Summary of commercial anti-human APOL1 antibody testing in Western blotting.

| | Western blotting | | |
|---|---|---|---|
| **anti-APOL1 Antibodies (see Table 1A for code)** | **dilutions used** | **detects APOL1 (40-42kDa doublet)** | **detects other bands** |
| 1A | 1:500-8000 | yes | 45kDa[1,5], 50-60kDa[1,2,4], 110kDa[2,4,5] |
| 1B | 1:500-8000 | yes | 45kDa[1], 50-60kDa[1,5], 110kDa[5] |
| 1C | 1:500-8000 | yes | |
| 1D | 1:500-8000 | yes | 50-60kDa[2,3] |
| 1E | 1:500-8000 | yes | 50-60kDa[4] |
| 1F | 1:500-8000 | yes | 50-60kDa[4] |
| 1G | 1:500-8000 | yes | 50-60kDa[3] |
| 2 | 1:1000-2000 | yes | |
| 3 | 1:1000-2000 | yes | 35kDa[1], 50-60kDa[1,2,4,5], 110kDa[1,2,5] |
| 4 | 1:1000-2000 | yes | 50-60kDa[4], 80kDa[4], 25kDa[3] |
| 5 | 1:1000-2000 | yes | 25kDa[3,4,5] |

[1]human proximal tubule cell lines
[2]human serum
[3]CHO cells
[4]human podocyte cell lines
[5]mouse whole kidney/glomerular extract

**Supplemental Table 1C.** Summary of commercial anti-human APOL1 antibody testing in human kidney tissue, and in mouse kidney tissue as a negative control.

| | | **Immunohistochemistry and/or Immunofluorescence** | | | | | | | |
|---|---|---|---|---|---|---|---|---|---|
| anti-APOL1 Antibodies | | Reactivity in human kidney | | | | Reactivity in normal mouse kidney | | | |
| (see Table 1A for code) | dilutions used | POD | ENDO | PEC | PT | POD | ENDO | PEC | PT |
| 1A | 1:200-400 | ++ | ++ | - | +++ | - | - | - | - |
| 1B | 1:200-400 | + | - | - | ++ | - | - | + | + |
| 1C | 1:200-400 | ++ | - | - | - | - | - | + | - |
| 1D | 1:200-400 | ++ | - | - | - | - | - | - | + |
| 1E | 1:200-400 | ++ | - | - | - | - | - | - | - |
| 1F | 1:200-400 | ++ | - | - | - | - | - | - | + |
| 1G | 1:200-400 | ++ | + | + | - | - | - | + | - |
| 2 | 1:50-1:100 | - | - | - | + | - | - | - | + |
| 3 | 1:50-1:100 | +++ | + | - | + | - | + | - | ++ |
| 4 | 1:50-1:100 | - | + | - | ++ | - | - | - | +++ |
| 5 | 1:50-1:100 | ++ | ++ | - | +++ | - | - | + | + |

POD: podocytes — negative: -
ENDO: endothelium, vasculature — weak: +
PEC: parietal epithelial cells — moderate: ++
PT: proximal tubule — strong: +++

SUPPLEMENTAL DETAILED METHODS:

**Western blot protocol**

Tissue or cell lysates (25µg) in Laemmli buffer were resolved by denaturing polyacrylamide gel electrophoresis using 4-20% gradient Tris-glycine gels followed by electrophoretic transfer to polyvinylidene difluoride membranes. Membranes were blocked with 5% nonfat dry milk in PBS with 0.2% Tween-20 at room temperature for 1hr and incubated with anti-APOL1 antibody (1:500 to 1:8000 dilution in 5% nonfat dry milk in PBS with 0.2% Tween-20) overnight at 4°C. Membranes were washed with shaking three times in PBS containing 0.2% Tween-20, 10min each wash, probed with secondary antibody (Horseradish peroxidase-conjugated goat anti-rabbit/mouse IgG) at 1:20,000 dilution for 1hr at room temperature, and washed three additional times with shaking in PBS containing 0.2% Tween-20. Immunoreactive proteins were identified using Supersignal West Pico Chemiluminescent Substrate per manufacturer's protocol (ThermoFisher Scientific).

**Immunohistochemistry/Immunofluorescence protocol**

Fixation conditions tested: Neutral-buffered formalin (10%, Sigma-Aldrich), HistoChoice MB (EMS), Zinc Formalin (Z-fix, EMS), Acetic methanol (50% methanol, 10% acetic acid in water).
Antigen retrieval methods tested: No antigen retrieval, EDTA Antigen Retrieval Buffer (pH 9.0, Abcam), Citrate Buffer Antigen Retrieval (pH 6.0, EMS), Trilogy (Sigma-Aldrich).

Fresh kidney tissue was placed in one of the fixatives listed above, at 4°C overnight. Following fixation, tissues were dehydrated and blocked in paraffin using standard procedures. Paraffin blocks were cut into 4µm sections and deparaffinized using Clear-Rite (Richard-Allen Scientific), except for Trilogy which is a combined deparaffinization and antigen retrieval process. Deparaffinized sections were processed with one of the antigen retrieval methods above, all using the standard protocols provided by the manufacturer.

Following antigen retrieval, sections processed for immunohistochemistry used an avidin-biotin amplification process with horseradish peroxidase secondary antibodies (ABC kit, Vector Labs). Kit recommended conditions were followed for all steps including blocking endogenous peroxidases. Detection was with DAB chromogen and sections were counterstained with hematoxylin. For immunofluorescence, sections were blocked using 5% normal goat serum in PBS with 0.02% Tween-20 using sufficient volume to cover the tissue section and incubated at room temperature for 1hr. Without letting the slides dry, slides were tipped to drain off blocking solution and primary antibodies were added, diluted in PBS with 1% normal goat serum (see Supplemental Table 1C for dilutions). Primary antibodies were incubated overnight at 4°C in a humidified chamber. Slides were submerged in wash buffer and washed three times (2 washes in PBS with 0.2% Tween-20, 1 wash in PBS), for 10min each wash. Secondary antibodies were diluted 1:400 (goat anti-rabbit or goat anti-mouse with appropriate fluorophore, Vector Labs) in 1% normal goat serum in PBS, and incubation was for 1hr at room temperature in a humidified chamber. Slides were submerged in wash buffer and washed three times (2 washes in PBS with 0.2% Tween-20, 1 wash in PBS) for 10min each wash. Slides were dipped 10 times in ultrapure water to removed salts and mounted in Vectashield anti-fade mounting media with DAPI (Vector Labs). Alternatively, dehydrating the slides and use of a hard-set anti-fade mounting medium was also successful.

SUMMARY: In Western blotting, most antibodies were able to detect APOL1, although several had a weak APOL1 signal with stronger signals for the non-specific bands. Most antibodies also detected additional proteins (gels bands that did not coincide with the molecular weight of APOL1) which differed depending on the cell source. Some of these non-specific gel bands could be eliminate with pretreatment of protein extracts with deglycosylating enzymes (not shown) suggesting epitope recognition was dependent on glycans. In kidney tissue, APOL1 detection was better with the inclusion of an antigen retrieval process, although this frequently also increased non-specific protein detection. Boiling or pressure cooker methods with citrate buffer were best, and crosslinking fixatives (formaldehyde-containing) performed better than precipitating fixatives (alcohol-containing). The overnight fixation in formalin was the superior fixation method.
